# Effective Fungal SynCom Assembly in Wheat Depends on Field Cooccurrence Patterns

**DOI:** 10.64898/2026.11.24.768850

**Authors:** Brianna K. Almeida, Mara DeMers, Christine V. Hawkes

**Affiliations:** Department of Plant and Microbial Biology, North Carolina State University, Raleigh North Carolina, USA

**Author notes:** Address correspondence to Christine V. Hawkes.

**Keywords:** Synthetic Communities, diversity, priority effects, drought, wheat

## Abstract

Plant-associated microbial communities can improve plant resilience to stress and as such are a promising avenue for the future of agricultural technologies to improve crop health. Ideally, synthetic microbial communities (SynComs) are simplified versions of real-world communities that can recapitulate their function. However, SynCom applications often have variable results, we hypothesize is due to the gap between the ideal and realized community. Understanding the mechanisms that support effective SynCom assembly is a key step to unlocking this resource. We investigated how fungal assembly mechanisms (priority, diversity) impacted wheat response to drought. The SynComs contained a single pioneer fungus applied first, followed by low diversity (3 fungi) or high diversity (9 fungi) community inoculations. Plants were then grown under drought or well-watered conditions. We measured plant phenotypic traits and realized fungal community. Realized communities contained a small set of the inoculated SynCom taxa and were not impacted by any treatment. To understand this outcome, we asked if fungal traits or field distributions could predict fungal persistence. We found that persistent SynCom taxa were those that were highly connected hubs in the field network, suggesting that fungi that cooccur and interact with many taxa are more likely to succeed in SynCom applications. Initial and realized SynCom composition influenced plant performance but not in a systematic way. This study highlights the importance of considering natural microbial communities during SynCom design to increase establishment and success of fungal taxa to improve crop stress tolerance.

**Importance:** Increasingly frequent and severe droughts pose a threat to our food systems, with cereal crop yield projected to decrease in the coming years. Although plant breeding will improve drought tolerance traits, additional management tools are needed. The agricultural industry is increasingly looking towards synthetic microbial communities (SynComs) for solutions. Here, we used SynComs with reduced microbial community complexity to study the ecological processes that impact microbiome assembly and host plant performance. Our SynComs were composed of foliar fungi which were previously shown to improve plant drought tolerance. We demonstrated that fungal SynCom persistence was best predicted by their associations in nature with persistent taxa more likely to serve as central network hubs. This study points to natural microbiomes as initial models for the development of successful SynComs with improved persistence and plant benefits in the future.

## Introduction

Increasingly frequent and severe drought events pose a threat to crop systems with cereal yields projected to decrease 10-20% by 2050 (Kraklow et al. 2025). Wheat constitutes 18% of global Calorie consumption (Kusunose et al. 2022) and studies have shown a 22% reduction of wheat yield under experimental drought stress (Chisaka et al. 2026), making it important to find solutions to maintain productivity. Crop drought tolerance has been improved through traditional breeding and gene editing methods, but adaptive genetic mechanisms will likely be inadequate to offset the impacts of climate change (Kusmec and Schnable 2024). Exploitation of the plant microbiome might complement the limits of genetic tools. Plant microbiomes can improve host nutrient acquisition and increase resilience to biotic and abiotic stresses (Liu & Brettel 2019, Trivedi et al. 2020, Rogriguez et al. 2008). For example, *Panicum virgatum* inoculated with individual foliar endophytic fungi had up to 450% less water loss in drought conditions (Giauque et al. 2019).

Microbial inoculants often target individual taxa that are known to improve plant stress tolerance or resilience (Salse et al. 2024). However, the effects of individual inoculants can vary with environmental conditions (Shayanthan et al. 2022). To help stabilize microbial effects in the face of environmental variation, synthetic microbial communities (SynComs) made up of multiple microbial taxa can be used to simulate the function and structure of whole microbial communities. For instance, maize inoculated with SynComs made of naturally occurring, highly abundant bacteria had lower leaf temperatures, compared to uninoculated maize, when under drought stress (Armanhi et al. 2021). Despite these advances, commercial SynComs frequently lead to inconsistent yield responses, indicating the need for further improvements (Shayanthan et al. 2022).

SynComs can be assembled using diverse approaches. A common method is to design SynComs to meet specific needs based on strain function in culture. For example, plants inoculated with bacterial consortia that are tolerant to desiccation have greater performance than plants with single-strain inoculants under drought conditions (Molina-Romero et al. 2017). However, strain traits do not always translate into plant benefits (Kibido et al. 2019, Bolin & Lau 2026). Alternatively, SynCom membership can prioritize strains that have already been associated with desirable plant responses or the production of secondary metabolites that generate specific phenotypes, such as enhanced plant growth and protection against pathogen infection (Martins et al. 2023). Yet another approach to SynComs is to represent the core microbiome to capture common coexisting taxa and mimic the function of natural microbial communities (Dolinsek et al. 2016, Toju et al. 2020). These approaches emphasize the traits of individual taxa or strains in the SynCom, but success tends to be unpredictable (Wang et al. 2025, Delgado Baquerizo et al. 2025). To improve SynCom performance, one possibility is to use ecological community principles that govern the function and stability of natural microbial communities.

Incorporating ecological community assembly mechanisms in SynCom design could improve SynCom effectiveness (Shayanthan et al. 2022). In particular, the order and timing of microbial colonization can influence community assembly and function as taxa that arrive first can affect later colonization. Early colonizers (hereafter “pioneers”) can inhibit the colonization of other microbes by taking up initial nutrients, occupying the available niche spaces or priming plant defense responses (Debray et al. 2022, Zamioudis & Pieterse 2011). Pioneers may alternatively facilitate colonization of subsequent taxa by changing plant growth and weakening the defense strategies of the host plant (Debray et al. 2022). For example, arrival order of fungal symbionts affects later pathogen colonization and disease in plants ranging from grasses to trees (Grunberg et al. 2023, Leopold & Busby 2020). Whether arrival order can more broadly alter microbial community assembly and function is a key gap in our understanding of SynComs.

Community diversity and richness can also impact SynCom performance. A hump-shaped diversity-function relationship is common in ecosystem ecology, with maximal function at intermediate levels of diversity (Tilman et al. 1997, Tilman et al. 2014, Jaillard et al. 2014). In microbiomes, we expect that functional redundancy will result in peak function before diversity is maximized. Beyond that optimum, competition between microbes may reduce function through compensatory competitive interactions (Tilman et al. 1996). However, microbiome studies have shown a variety of diversity-function trends. Some studies support this hump-shaped hypothesis (Almeida et al. 2024). Other studies show both positive and negative linear effects (Liu et al. 2023, Aponte Rolon et al 2024). For example, increasing bacterial strain diversity was associated with decreased disease control in pepper (Lu et al. 2026). In contrast, a meta-analysis of 97 SynCom studies showed that as taxon richness increased, so did inoculation effects on the host (e.g., plant biomass, yield, seed germination, etc.) (Liu et al. 2023). Regardless of the exact form of the relationship, these results suggest that microbial diversity plays an important role in microbiome-mediated plant phenotypes.

We addressed the intersection of priority and diversity in SynCom development to understand whether these ecological principles could drive the success of fungal phyllosphere inoculants and their resulting impact on plant stress phenotypes. To do this, we tested 30 fungal SynCom treatments on wheat response to drought. When developing the SynCom treatments we considered an assembly mechanism (priority) and a community characteristic (diversity) by manipulating order of arrival with pioneers introduced before subsequent communities consisting of 3 or 10 fungal taxa. We also created variation in each diversity level by generating replicate compositions that differed in phylogenetic distance.

We tested the following four hypotheses. 1) We expected that pioneers would affect colonization success low diversity communities would be especially affected because they lack functional redundancy. This may be the result of multiple mechanisms, such as occupying space or priming the host immune system. 2) We hypothesized that successful fungal colonizers from SynComs would be taxa that were abundant and highly connected in field communities on wheat because these are likely able to interact with many other fungi (Agler et al. 2016). 3) We further expected drought to favor more drought-tolerant fungal taxa, where wheat under drought might select for more phylogenetically diverse and larger communities if they contain more potential pathways to ameliorate host drought stress (Rodriguez et al. 2003, Maherali & Klironomos 2007). 4) Assuming that SynComs affect plant performance, we expected a range of responses where some SynCom treatments would ameliorate drought stress and others might exacerbate it given that many endophytes are latent or facultative pathogens (Rodriguez et al. 2009, Indermaur & Westrick 2026).

## Materials and Methods

### Experimental design

To test priority and diversity effects on how the wheat microbiome assembles and impacts plant drought responses, we grew wheat with synthetic fungal communities that contained a single pioneer fungus applied first, followed by low diversity (3 fungi) or high diversity (9 fungi) community inoculations. To maximize community variation, each level of diversity included 3 levels of phylogenetic relatedness (low, medium, high). Plants were grown in sterile microcosms where they were paired, with one plant assigned to well-watered conditions and the other to drought. Treatments were replicated 6 times, including controls, for a total of 382 microcosms. Treatment details are provided below.

### Fungal collection and consortia selection

Cereal leaves of varying ages were collected in and around wheat fields at 7 sites across North Carolina USA in March and April 2022 (Supplemental Figure 1). These sites varied in their climatic conditions and soil properties. To isolate fungal cultures, leaves were surface sterilized by successive washes with tap water for 30 s, 95% ethanol for 30 s, 0.5% sodium hypochlorite solution for 2 min, 70% ethanol for 2 min, and 1 min in sterile water. Surface-sterilized leaf fragments were placed in 1.5 mL slant tubes containing potato dextrose agar (PDA) and allowed to grow at room temperature (22°C). Individual fungal isolates were then transferred from slant tubes to PDA plates with 50 µg/mL ampicillin and gentamycin resulting in 2823 total isolates, from which we selected 95 to move forward that varied in phylogeny, based on morphotyping.

To identify these isolates, we extracted fungal DNA using the E.Z.N.A. HP Plant & Fungal DNA Kit, and PCR-amplified and sequenced the following DNA regions: internal transcribed spacer (ITS), large subunit (LSU), beta tubulin (Btub), actin (ACT), and translation elongation factor-1α (TEF1) (Koehler at al. 2019). Sanger sequencing took place at the North Carolina State University Genomic Sciences Laboratory (NCSU GSL). Sequenced regions were concatenated for each isolate and aligned using MAFFT (UniPro Ugene) to reference sequences acquired from the NCBI database (Katoh et al. 2002). To remove poorly aligned sections of the sequences, we used Gblocks (Talavera et al. 2007, Castresana 2000). We created a maximum likelihood tree using RAxML 8 based on the alignments (Supplemental Figure 2; Stamatakis 2006, Stamakis 2014). From this tree, we selected 42 isolates that varied in phylogenetic relatedness and were not closely related to foliar pathogens in the NCBI database.

From the 42 fungi, we selected five that spanned the phylogeny of isolates that would serve as “pioneers” in our growth chamber experiment to test fungal priority effects. The pioneers and the remaining 37 fungi were used to create SynCom treatments that varied in the number of isolates as well as the phylogenetic relatedness of each SynCom (Figure 1). SynComs contained 4 isolates or 10 isolates (including the pioneer isolate) to create “low” and “high” diversity treatments, respectively. Within each diversity treatment, we included communities with three levels of phylogenetic relatedness (“low”, “medium”, and “high”) for a total of 30 different consortia (Figure 1) (R packages *phytools (2.5-2), ape(ver.5.8-1)* Revell 2024, Paradis & Schliep 2019). Each SynCom treatment was applied to 6 replicate microcosms. In addition to these treatments, a pioneer monoculture inoculation treatment was created to understand how the different pioneer isolates impacted plant performance without additional isolates. There were also 6 microcosms that received a control treatment of sterile water.

**Figure 1:**
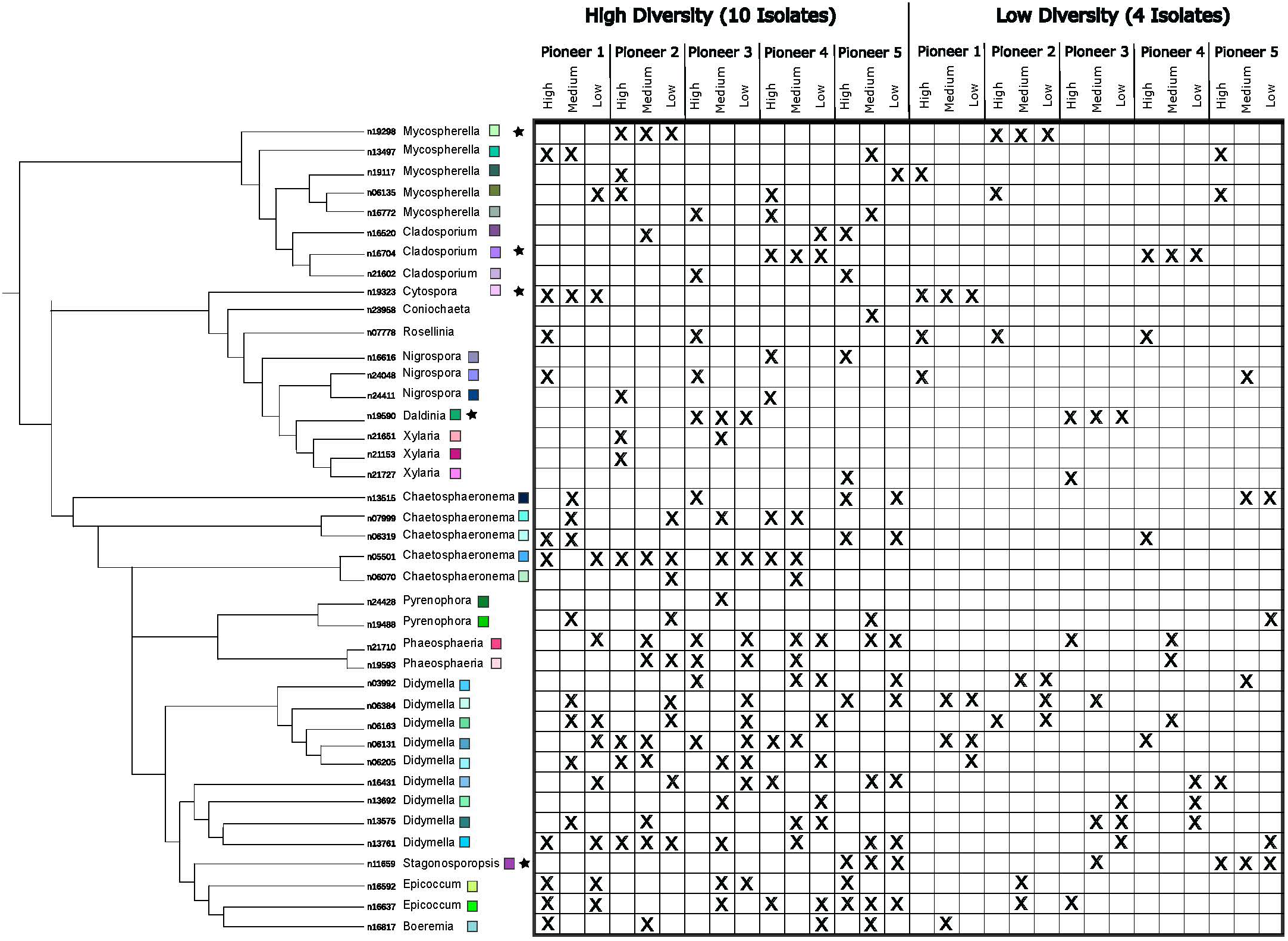
The phylogenetic tree shows Syn Com membership and relationships between inoculated taxa based on the multilocus RAxML 8 maximum likelihood tree. This is a modified version of the phylogenetic tree of Supplemental Figure 2, only maintaining strains used as inoculants in the experiment. Stars indicate isolates that were designated as pioneers. Colored squares match the colors in Figure 3. SynCom treatments are listed across the top, with X’s in the columns indicating isolates that were included in each treatment. Pioneer 1 is *Cytospora*- 19323, Pioneer 2 is *Mycosphaerella*-19298. Pioneer 3 is *Daldinia*-19590. Pioneer 4 is *Cladosporium*-16704. Pioneer 5 is *Stagonosporopsis*-11659.

### Seed preparation and germination

Our focal plant was the soft red winter wheat (*Triticum aestivum*) cultivar AGS2000 (Johnson et al. 2002). It is adapted to growth in the southeastern United States, is only moderately resistant to common wheat diseases, and performed well in microcosms in our previous work. Wheat seeds were sourced from US Department of Agriculture Agricultural Research Service fields in North Carolina. To remove any existing seed endophytes, wheat seeds were scarified with fine grit 220 sandpaper, submerged and shaken in a series of washes (95% ethanol for 30 s, 8.25% sodium hypochlorite and 0.01% Tween 20 for 10 min, 95% ethanol for 1 min, and sterile water × 3), and allowed to imbibe for 1 h in sterile water with 4% 500 ppm gibberellic acid (GA). Seeds were then germinated in 24-well plates containing MS agar, 350 µL of 500 ppm GA, and 300 µL of 1% chlorothalonil fungicide (Daconil). After 10 days, visibly uncontaminated germinants were transferred to 267 mL pots that contained sterilized montmorillonite clay (e.g., Turface, Oil-Dri, Pro’s Choice) that was autoclaved three times for 1 h with 24 h rest in between.

### Fungal inoculations

Plants were first inoculated with one of the five pioneer isolates or a sterile water sham inoculum. To create the fungal inocula, we grew fungi in 1/4 strength potato dextrose broth (PDB) for 10 days. We centrifuged the liquid media at 5,000 × g for 10 min, removed the supernatant, and rinsed the fungi with sterile water. Fungal isolates were homogenized using a blender (Duralast Classic, Oster). Hyphal fragment concentration was diluted to 1 × 10^4^ hyphal fragments mL^-1^. The pioneer inoculants were then loaded into an airbrush (Paasche H Siphon Feed Airbrush) and sprayed on the wheat plants (approximately 1 mL per plant). Airbrush parts were washed and sterilized with 8.25% sodium hypochlorite and 95% ethanol between isolates. After 2 weeks of plant growth, the SynCom treatments were sprayed onto the plants (Figure 1). These treatments were created similarly to the pioneer inoculate and were assembled by diluting all fungal isolates to 1 × 10^4^ hyphal fragments mL^-1^. Isolates for each SynCom were mixed in equal volumes and ∼1 mL was applied to each plant. At the time of community addition, control microcosms also received another sham inoculum of sterile water. Due to space limitations in the growth chamber, replicates for each treatment were divided into two time blocks in a stratified random design to ensure equal treatment representation in each block.

### Microcosms and growth chamber conditions

Plants were grown in pairs in microcosms comprised of sterile 13 L clear polyethylene bags (Twirl ‘Em, Labplas, Quebec, Canada). These were modified with a tube system for watering, where a 5-cm vinyl tube was put through the clear polyethylene bag and sealed with a luer lock ring and thread and covered with parafilm, and an 8 cm^2^ window of 15-µm nylon mesh designed to allow for gas exchange while minimizing contamination. Both pots in a microcosm received the same fungal treatment. The microcosms were kept in a walk-in growth chamber in the NCSU Phytotron with a 12-hour day/night cycle at 24/16°C. Microcosm positions were randomized weekly. For the 7 weeks prior to drought treatment implementation, pots were watered with approximately 60 mL/week of quarter strength 15-5-15 NPK fertilizer (Jack’s Professional, JR Peters, Inc., Allentown, PA, USA).

### Drought application and harvest

Two weeks after consortia inoculation, pots in each microcosm were randomly assigned to drought or well-watered treatments. Water was withheld from plants in the drought treatment while plants in the well-watered treatment received 60 mL of sterile water each week. Drought conditions were maintained for approximately 3 weeks until the majority of drought treatment plants showed signs of stress (wilting and leaf tip browning). Prior to harvest, we measured height, leaf dieback, stomatal conductance (LI-600, LiCor, Lincoln, NE), and severity of disease symptoms if there were any lesions present on the plants. Above and below ground tissues were then collected and separated. We measured aboveground fresh weight, then selected 2-3 leaves which were weighed, flash frozen in liquid N2, and stored at −80°C for DNA extraction. Remaining biomass was dried at 60°C for 15 days to obtain total dry biomass.

### Amplicon library preparation

Final realized fungal communities in wheat leaves were characterized with ITS amplicon sequencing. Frozen leaf tissue samples were lyophilized for 2 days and ground to powder (HG-600 Geno/Grinder 2010 tissue homogenizer, Cole-Parma SamplePrep) DNA was extracted from 10 mg of leaf tissue using the Mag-Bind Plant DNA DS Kit (Omega Bio-tek, Inc). We amplified the ITS1 region using the ITS-F_KYO2 and ITS86R primers (Toju et al. 2012, Scibetta et al. 2020). PCR consisted of 12.5 µl of KAPA HiFi HS ReadyMix (Roche Diagnostics, Indianapolis, IN), 0.5 µl of each primer (10uM), 8.85 µl molecular water, 2.5 µl of DNA (around 20 ng/µl). We also spiked each PCR with three fungal synthetic sequences (0.25 fg) (Palmer et al. 2018; Supplemental Table 1 and 2 with primers and internal standard sequences sequences). Illumina libraries were prepared following a modified version of the manufacturer’s workflow (Illumina 2013 /Modified protocol in supplement). Sixteen water negative controls with eight DNA extraction controls and eight PCR controls and 382 DNA samples were submitted to the NCSU GSL for amplicon sequencing on the Illumina MiSeq platform (v2 250 bp paired end).

### Bioinformatics

To process the amplicon sequences, first, we filtered low-quality samples with *fastqc* (sample quality < 21, Andrews 2010) and removed adapters and primers using *TrimGalore* and *cutadapt* (Krueger 2019, Martin 2011). Low-quality base pairs were trimmed using *filtandtrim* from the *dada2* package (v. 1.34.0 e.g., max EE=2, trunQ=2, rm.phix=T, Callahan et al. 2016). We used *dada* to infer amplicon sequence variants (ASVs) and remove chimeras (Callahan et al. 2016) following default package conditions (method=” consensus”). ASVs were curated with *MuMu* (Mahe 2024), and contaminants were removed (R package *decontam* v. 1.26.0; Davis et al. 2017) based on ASV frequencies in the negative controls.

To match ASVs to the original isolates based on their ITS sequences, we used *blastn* (Altschul et al. 1990) with 80% coverage and 84% identity. ASVs that did not have a match were placed on a fungal phylogenetic reference tree using the tree-based alignment selector toolkit (T-BAS, Carbone & White 2025). Based on cophenetic distances (R package *ape*), ASVs with a distance less than 0.2 were merged and then compared to reference sequences using a MAFFT multiple sequence alignment. The remaining unmatched ASVs were identified as their closest reference taxa in the T-BAS tree and were considered environmental contaminants. To further understand contaminant taxa, ASVs that were merged were put into NCBI BLAST to understand their possible fungal community impacts.

### Fungal Trait Assays

To determine if fungal traits related to drought tolerance could be used to predict consortia effects on the plant host, we assayed several fungal traits in culture independent of the plant for each of the 42 fungi. Specifically, we measured one trait related to plant growth promotion (indole acetic acid (IAA) production) and two traits related to fungal stress tolerance (desiccation recovery and osmotic sensitivity) that we hypothesized would be relevant to their effect on host performance. To quantify fungal IAA production, we used a colorimetric assay based on Ikram et al. 2022. Color was allowed to develop for 10 minutes after which we measured the optical density at 540 nm using a plate reader (Tecan Infinite M Plex Multi mod plate reader). To assess fungal recovery, we compared fungal growth in 6-well plates in which 3 wells were sealed with a permeable membrane (Breathe Easier, Diversified Biotech) to allow evaporation, and 3 wells were sealed with an impermeable membrane (Parafilm) to prevent water loss. Plates were incubated at 28°C and after 7 weeks, we re-wet all wells with 300 µL of sterile water and plated isolates on new PDA plates and waited for recovery. To determine osmotic sensitivity, fungi were grown in M9 liquid four levels of NaCl (0, 40, 80, 120 gL^-1^ NaCl; Bell & Gonzalez 2009) and fungal biomass was measured after 7 weeks. Osmotic sensitivity was calculated as the linear slope of growth across the NaCl gradient.

### Statistical Analyses

To address the effects of pioneers, diversity, and drought, we performed a multivariate analysis of variance (MANOVA) using a residual randomized permutation procedure (RRPP) to analyze the realized fungal community composition as a function of the initial SynCom treatments (fixed effect), drought treatment nested in microcosm (random effect), and time block (random effect) (R package *RRPP* v. 2.1.2; Collyer and Adams 2018). SynComs have a naturally low amplicon diversity and a lower overall sequence yield and low confidence in taxa abundances (Illumina 2021). Therefore, all tests that include the realized fungal community are based on the presence/absence of taxa in the community. We performed a post-hoc analysis using *pairwise* (R package *RRPP* v. 2.1.2). We also tested whether dispersion between community composition varied with fungal treatment using *betadisper* (R package *vegan* v. 2.7-1). Results were visualized using an NMDS ordination (Figure 2).

**Figure 2:**
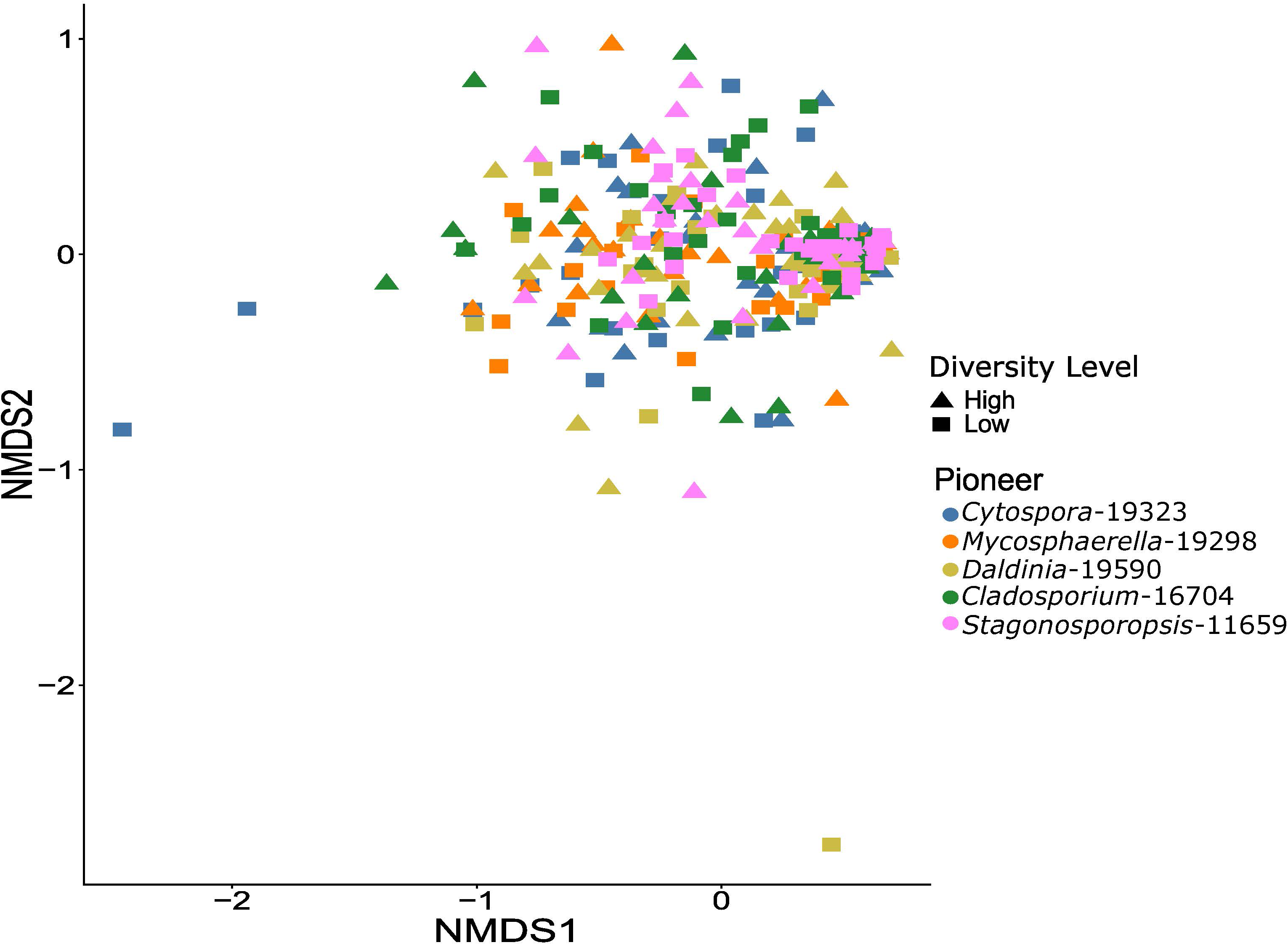
Nonmetric multidimensional scaling of realized fungal communities including contaminants. Each point is the community of one leaf sample colored by pioneer treatment with the shape indicating the diversity level treatment.

To better understand how field distributions could predict the persistence of inoculated fungal taxa, we examined the abundance and connectivity of experimental fungi in the field. For this investigation, we define persistence as a fungal taxon occurring in at least one sample. To understand field distributions, we queried a database of wheat sequencing data collected the same 7 wheat fields in North Carolina from which the cultures were isolated (Supplemental Methods). We investigated the cooccurrence of fungi in the field that persisted in the experiment using network analyses, specifically using SparCC to estimate correlations between taxa (Friedman & Alm 2012, R Package NetCoMi; Peschel et al. 2020). The field network was visualized, and network qualities were determined with Cytoscape (Shannon et al. 2003). We performed a MANOVA with persistence as an explanatory variable (persistent, not persistent, or Untargeted ASVs as categories) and degree, radiality, closeness centrality, and betweenness centrality as response variables. In addition, we performed a MANOVA using a residual randomized permutation procedure (RRPP) with persistence category and relative abundance of taxa in the field community.

To determine the relationship between SynComs and plant performance, we analyzed plant responses to both the inoculated and realized communities. We first performed a MANOVA on plant size and growth metrics that were measured at harvest (shoot mass, root mass, height, root-shoot ratio, leaf dieback, and ratio of leaves with lesions to total leaf number) as a function of fungal SynCom treatment and water treatment (fixed effects) and microcosm and time block as random variables (*lmer* R package *lme4* v. 1.1-37, Bates et al. 2015). We performed a separate ANOVA with leaf-level stomatal conductance as a response variable, because this was measured at a different time from the harvest. Before analyses, plant performance metrics were assessed for normality using Shapiro-Wilk tests and transformed as appropriate for each metric.

To determine if realized fungal community composition was associated with any plant performance metrics, we performed another RRPP with realized fungal community as a Jaccard distance matrix as the dependent variable and plant performance metrics as the explanatory variables. Because realized community composition was associated with some plant performance metrics, we dug further into those results to identify key taxa associated with performance outcomes. We investigated which isolates influenced plant performance based on their presence/absence in the realized community composition with random forest models including the interaction with water treatment in the model (fixed effect) (R package *MaAsLin 3* v. 1.1.2; Nickols et al. 2026). To examine whether fungal trait data could be used to predict plant performance metrics, we used a linear mixed model with the average community IAA concentration, average osmotic sensitivity, and desiccation recovery as explanatory variables. All analyses were conducted using R ver 4.4.1 (R Core Team 2024).

## Results

### SynCom Realized Composition

There were a total of 738 ASVs after removal of wheat and contaminant reads in all treatments. On average, 7 ASVs were assigned to each experimental isolate and 44 ASVs were assigned to contaminants. All samples had some level of contamination either from seed borne fungi or from the environment of the growth chamber (Supplemental Figure 3). Due to their ubiquity in the fungal communities at harvest, they were included in all subsequent sequenced community analyses. Realized fungal community composition was associated with initial SynCom treatment (*P* < 0.001, *R*^2^ = 0.11, Figure 2), but not with drought (Table 1). We used planned comparisons to determine if community characteristics, pioneer identity and fungal diversity impacted realized fungal community. We found SynComs with two pioneers (*P*<0.001, *R*^2^ = 0.023) differed in realized community composition, specifically those with pioneers *Cladosporium*-16704 and *Stagonosporopsis*-11659 (Supplemental Table 3).

**Table 1:** PERMANOVA (RRPP) results for realized community composition based on Jaccard dissimilarity and experimental treatments.

|  | df | SS | MS | Rsq | F-value | Z | P-value |
|---|---|---|---|---|---|---|---|
| Initial SynCom Treatment | 30 | 13.699 | 0.456 | 0.117 | 1.59 | 5.66 | <0.001 |
| Microcosm | 1 | 0.488 | 0.488 | 0.004 | 1.4 | 1.32 | 0.093 |
| Consortia: Microcosm | 30 | 9.89 | 0.329 | 0.085 | 0.994 | -0.058 | 0.510 |
| Consortia: Microcosm: Drought | 31 | 8.92 | 0.288 | 0.076 | 0.867 | -2.58 | 0.996 |
| Residuals | 208 | 69.05 | 0.332 | 0.592 |  |  |  |
| Total | 300 | 116.737 |  |  |  |  |  |

### Persistence of SynCom Taxa

Of the 42 fungal isolates included in the SynCom treatments, only 13 (31%) were recovered at the end of the experiment based on amplicon sequencing of the leaf tissue. These 13 isolates included two of the five pioneers (Figure 3). All high diversity communities had at least one isolate remaining from the initial community inoculum. This was not the case for the low diversity communities, where nine of 15 whole communities did not persist and were instead comprised solely of contaminants (Supplemental Fig S3).

**Figure 3:**
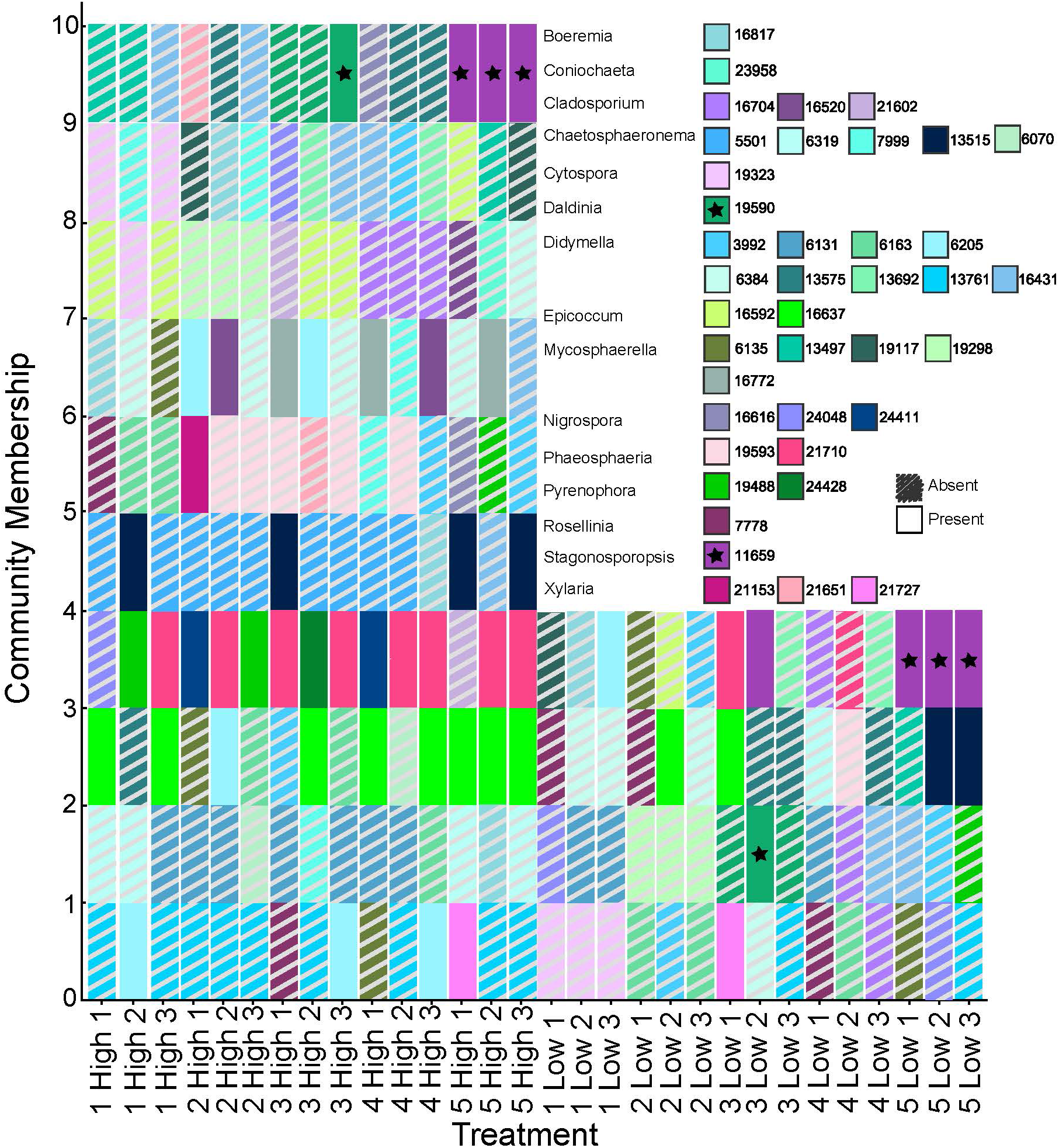
SynCom membership and isolate persistence. Each bar is an isolate that was included in the inoculum for each initial SynCom treatment. Bars are colored by isolate and hashed lines indicate that the isolate was absent from the realized community based on amplicon sequencing. Stars indicate the pioneer taxa that persisted to the end of the experiment.

To better understand why some taxa persisted, we examined the distributions of inoculated taxa in field community networks. This was accomplished by amplicon sequencing foliar fungal wheat communities from the same fields where fungi were isolated, using leaves collected at the same time. We found 16 of the 42 experimental isolates in the field data, including 12 of the 13 persistent in the growth chamber experiment. Persistent taxa were important nodes in the field community network (Figure 4A) that were 122% more connected (degree, *P=* 0.009), 15% more central (closeness centrality, *P*=0.54), and 1% more proximal to all other nodes (radiality, *P*=0.011) compared to non-persistent taxa from the experiment (Table 2). One of the pioneers, *Stagonosporopsis*-11659, was persistent in the experiment and central to the network. In addition, relative abundance of SynCom taxa in the field was not associated with persistence in the experiment (*P=*0.298, Figure 4B, Supplemental Table 4). The remaining three pioneers and other inoculated fungi did not appear in the field data.

**Figure 4:**
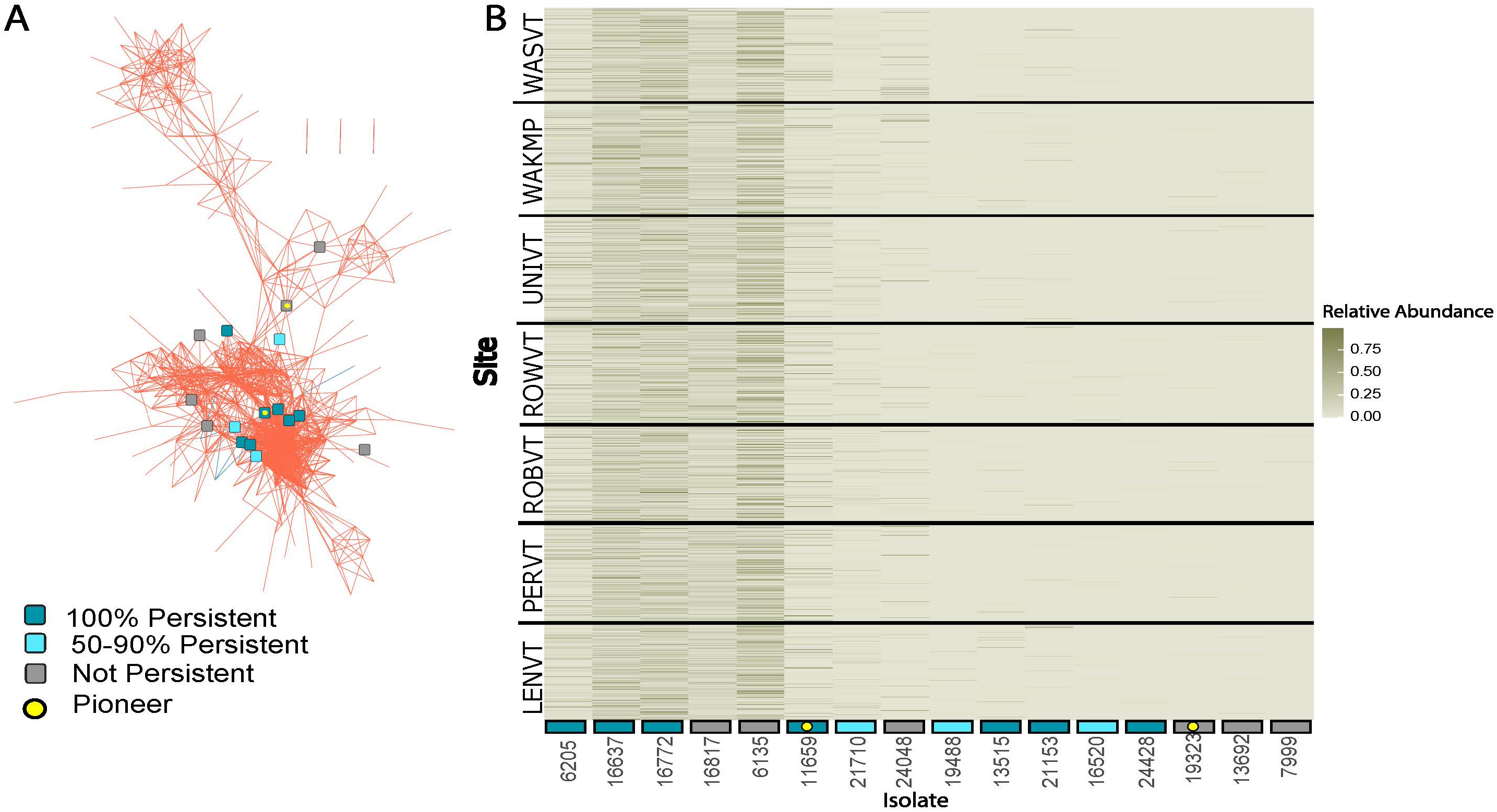
Taxa that were persistent in the SynCom experiment are largely central to field community networks. Dark cyan boxes represent taxa that were persistent across all communities where they were inoculated. Light aqua boxes represent taxa that were 50-90% persistent and gray nodes are taxa that were not persistent. Yellow dots indicate pioneer taxa. (a) Fungal association network with nodes at the intersections of edges for all other taxa from the field removed for figure clarity. (b) Heat map of the relative abundances of SynCom taxa in the field, with persistence indicated by color.

**Table 2:** Network statistics (average ± 1 standard deviation) for inoculated SynCom taxa that were persistent or not persistent, as well as untargeted taxa from the field community. Degree is the number of connections to other nodes. Radiality describes the organization of nodes into hub and spoke structures. Betweenness centrality is a measure of how frequently a node appears on the shortest path between other nodes. Closeness centrality is the ratio of 1/ the sum of paths from the node of interest to all other nodes.

|  | Untargeted ASVs | SynCom ASVs<br>Not-Persistent | SynCom ASVs<br>Persistent |
|---|---|---|---|
| Degree | 11.5±10.41 | 10.33±8.33 | 21.5±12.11 |
| Radiality | 0.951±0.016 | 0.958±0.008 | 0.966±0.005 |
| Betweenness Centrality | 0.0099±0.026 | 0.014±0.014 | 0.024±0.042 |
| Closeness Centrality | 0.282±0.13 | 0.292±0.039 | 0.327±0.031 |

### Plant Performance

Plants were smaller in the drought treatment (Supplemental Table 5), with 11% less shoot mass and 18% less root mass compared to well-watered plants (Supplemental Figure 4). Regardless of drought, the initial SynCom treatments affected plant height, root mass, and root-to-shoot biomass ratio (Supplemental Table 5). SynComs generated 11% variation in plant height and 49% in root mass (coefficient of variation; Figure 5). Plant performance metrics did not depend on inoculated pioneer identity or treatment diversity but were weakly associated (R^2^ < 0.01) with realized fungal community composition (Supplemental Table 6).

**Figure 5:**
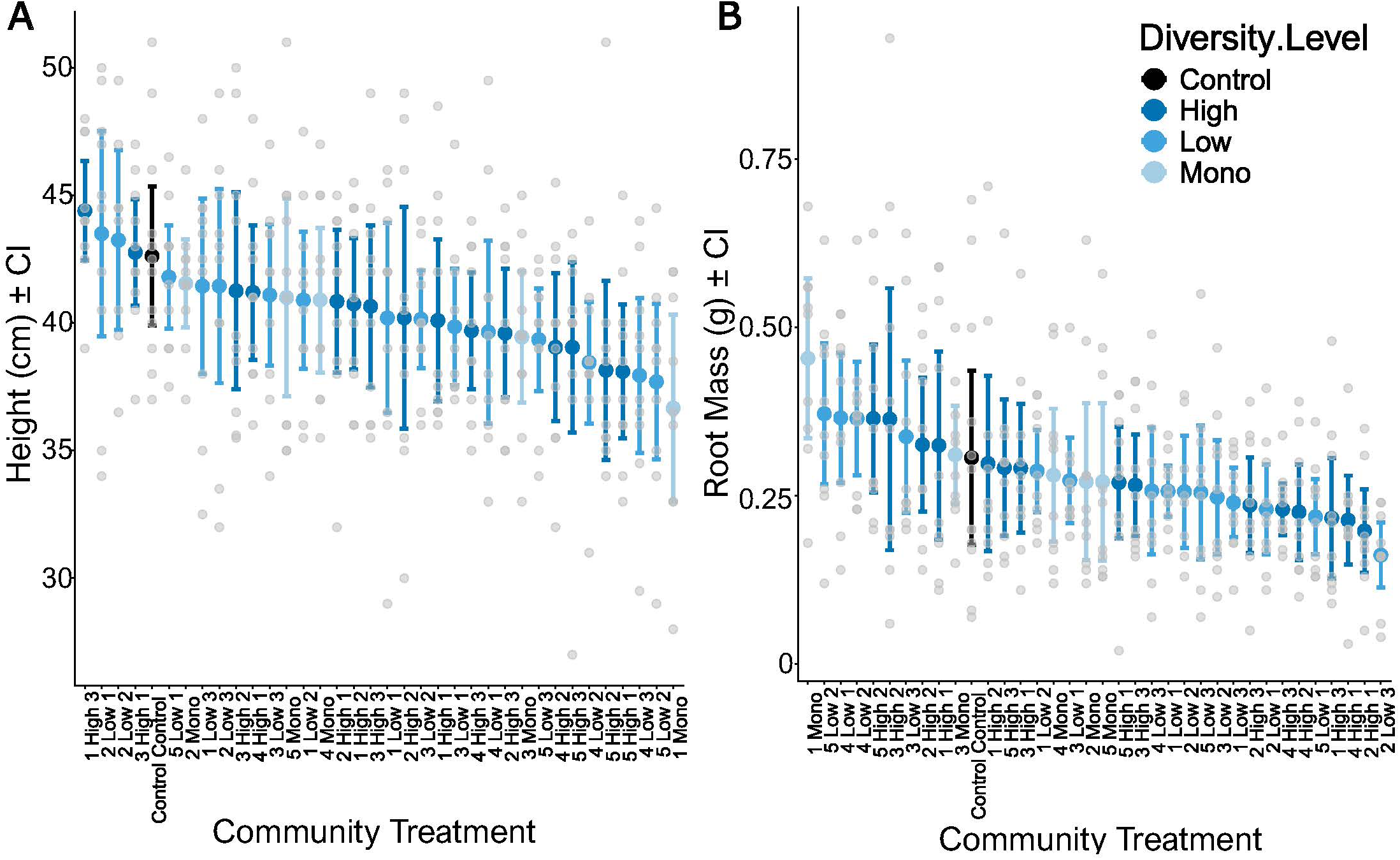
Initial synthetic community inoculum was associated with (a) plant height (*P*=0.015) and (b) root mass (*P*=0.0029). Dark blue indicates communities with high diversity (10 taxa). Medium blue indicates communities with low diversity (4 taxa) and light blue indicates communities with a single pioneer inoculant. Black indicates the control treatment which included an inoculum with sterile water. The light gray dots indicate individual plant measurements. Points with bars indicate mean ± CI.

We tested whether the limited effects of realized SynCom fungal community composition on plant performance were due to only a subset of taxa potentially affecting the plant host. Based on a random forest model with isolate presence as the predictor (including contaminants) and each plant performance metric as the responses, we found that plants were 3.3% shorter in the presence of the pioneer *Stagonosporopsis*-11659 (*P*=0.02), the inoculant *Pyrenophora*-24428 (*P*=0.012), and the contaminant *Fusarium* (*P*=0.033). There was also 9% less root mass in the presence of the contaminant *Elsinoё* (*P*=0.029). In addition, the presence of *Didymella*-6205 (*P*=0.023) and the contaminant *Fusarium* (*P*=0.040) were associated with a 51% increase in the proportion of diseased leaves (Figure 6).

**Figure 6:**
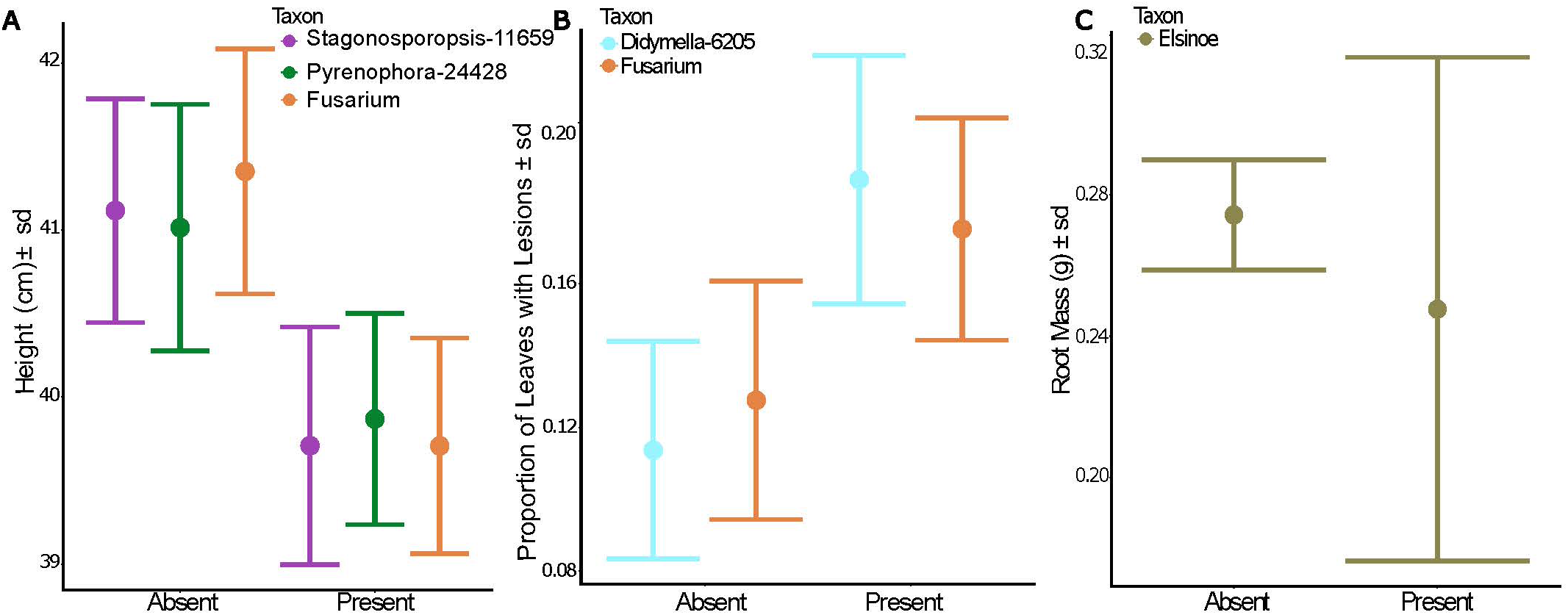
Presence of some taxa in the realized fungal communities influenced plant performance metrics including (a) height, (b) leaf lesions, and (c) root mass. Points with bars indicate mean ± SD. Colors for each fungus reflect Figures 1, and 3.

When investigating whether fungal traits are predictive of plant performance, we found that average osmotic sensitivity of each community was significantly associated with shoot mass and root mass, however, very weakly. (Supplemental Table 7, Supplemental Figure 5 R^2^ < 0.04). If osmotic sensitivity and IAA are averaged based on realized community composition, then IAA production is associated with height, leaf lesions, and the proportion of brown leaves, but again they were weakly associated (Supplemental Table 8, Supplemental Figure 5, R^2^ <0.05). In addition, trait dissimilarity, based on Euclidean distance of each isolate within the realized community composition, was associated with plant performance traits, specifically height leaf (p=0.02, R^2^=0.02), dieback (p=0.006, R^2^=0.02), and leaf lesions (p<0.001, R^2^=0.05). Desiccation recovery was not analyzed because only two isolates failed to recover, *Cytospora*-19323 and *Rosellinia*-7778. Both isolates were not persistent in the growth chamber experiment, and only the *Cytospora* was present in the field network. This *Cytospora* was not central to the network and had a low number of connections.

## Discussion

Realized communities contained only a small subset of the applied SynComs, with nine of the 42 inoculated taxa detected at the end of the experiment. These communities were only loosely dependent on arrival order and not at all on diversity or drought. Instead, persistent taxa from the growth chamber experiment were central to the field community network, suggesting that highly connected taxa in field communities are more likely to successfully establish in SynComs. Although the SynComs resulted in a range of plant sizes consistent with our hypotheses, these results were poorly predicted by either ecological assembly mechanisms or trait distances of either the original or realized communities. Based on these results, SynCom development may be improved by considering existing patterns of cooccurrence in natural communities to define possible species pools. Using these cooccurrence patterns as a first step in the design process of SynComs may increase establishment success of taxa selected from that pool based on other considerations.

Well-connected taxa (i.e., “keystones”) that can facilitate the establishment, persistence, and function of other taxa are likely to be an important aspect of successful establishing SynComs. Outside of highly connected fungi, we had very low taxa persistence in this experiment. In such low diversity, low abundance, stressful, microbial systems like that of leaves, cooperative interactions that were missing here may be required for establishment. Previous studies have shown that the inclusion of helper taxa that synergize with keystones could increase community stability and functional performance (Wang et al. 2025, Xun et al. 2021).

Competitive and antagonistic interactions can also occur within simplified communities like SynComs particularly when resources are scarce, preventing persistence of some community members (Chaudry et al. 2020, Wang et al. 2025, Westhoff et al. 2021). The broader use of keystone microbial taxa in SynComs could be tested by examining how SynCom results from previous studies relate to field distributions based on public databases. It is unclear, for example, whether the same rules that apply to low diversity leaf habitats would enable successful establishment of SynCom taxa in high diversity, high abundance habitats such as soil, where competition might be more intense (Xun et al. 2026).

Of the persistent taxa, only two of the five pioneers were detected in realized communities, specifically *Daldinia*-19590 and *Stagonosporopsis*-11659, which may indicate their ecological roles influencing community composition. *Stagonosporopsis*-11659 was in six of its seven SynComs and one of the most connected nodes in the field community network. It also persisted in the one community where it did not serve as a pioneer. Members of the genus *Stagonosporopsis* are well known pathogens of cucurbits (Zhang et al. 2026) but are beginning to be recognized as a mutualist including the recognition that they can shift between these conditions (Wei et al. 2024). In contrast, *Daldinia*-19590 only persisted in two of six SynComs and was not detected in the field data. This is unexpected as *Daldinia* can act as biocontrols and plant growth promoters that benefit their plant hosts (Athinuwat et al. 2026, Liarzi et al. 2016). An additional inoculated pioneer, *Cytospora*-19323, was detected in the field as a poorly connected node but did not persist in the experiment. This pioneer is often cited as a pathogen of a wide range of hardwood and coniferous trees, which may explain its poorly connected role in wheat field communities (Lin et al. 2023, Pan et al. 2020). The final two pioneers were not detected in either realized or field communities, suggesting a more transient lifestyle and limited presence on adult wheat plants (since they were able to be cultured). Host developmental shifts can impact transient microbes, and fluctuating habitats can limit establishment and survival over time, especially in low diversity leaf communities (Chaudry et al. 2020, Suteau et al. 2025). Because pioneer isolates were cultured from plants four weeks older than those used in this experiment, they may not have been well adapted to the developmental stage at which they were inoculated. Finally, it is possible that some pioneers were not compatible with the host genotype we chose for the experiment as fungi in the culture collection were isolated from field wheat of different cultivars (Leopold & Busby 2020).

Two pioneers influenced realized community composition, including one persistent (*Stagonosporopsis*-11659) and one transient (*Cladosporium*-16704). Although we might interpret this as an indicator that persistence is not required if the goal is to facilitate assembly of other taxa, our results suggest otherwise. In low diversity treatments, the transient pioneer was in communities in which no isolates persisted while the persistent pioneer resulted in one or two persistent taxa in realized communities. In high diversity treatments, the transient pioneer resulted in one, three, or four taxa persisting while the persistent pioneer was associated with consistent, persistent four-species communities. Previously, species of *Cladosporium* have been described as able to induce plant host defense response (Cai et al. 2007, Laugé et al. 2001) and secrete antifungal metabolites against other plant pathogens (Wang et al. 2013). It is possible that although transient, our *Cladosporium* pioneer was able to impact the subsequent community composition by priming plant defenses or altering leaf environments to be hostile to later arrivals. In addition, transient taxa, in general, can induce alternative stable states in microbial communities, for example, by changing the substrate pH, limiting the survival of certain taxa (Liu et al. 2024, Amor et al. 2020).

Inoculated pioneers may not have been the first to arrive based on observed fungal contaminants, and those already established taxa could have prevented the success of new arrivals (Supplemental Figure 3). The fungal contaminants are only contaminants in the sense that we did not inoculate them, as they were likely seed endophytes based on overlap of taxa with wheat seed studies (Clear & Patrick 1993, Becker & Cubeta 2026). Seed contaminants included a mixture of known pathogenic fungi and potentially beneficial endophytes. Of these, almost all treatments included *Parastaganospora nodorum*, a common necrotrophic wheat pathogen and the causal agent of septoria nodorum blotch (Adhikari et al. 2023, Richards et al. 2019), and multiple species of *Fusarium*, including the causal agents of *Fusarium* crown rot and members of the pathogenic *Fusarium fujikuroi* species complex (Qiu et al. 2020, Manghwar et al. 2021, Hamza et al. 2023, Ma et al. 2025, Thao et al. 2025, Hao et al. 2026). A few other taxa are known from other wheat studies but have not been tested for function with regards to wheat performance (Salamon et al. 2025). There were also some beneficial fungi that were consistently present, including *Aspergillus fumigatus,* which can prime wheat and maize and improve plant growth when exposed to stressors such as drought and salinity stress by promoting plant antioxidants and enzyme activity (Yassin et al. 2024, George et al. 2024). However, we cannot confirm if the contaminants were truly seedborne because seed microbiomes were not sequenced as part of this experiment.

Previous pioneers used in SynComs have been found to influence community change, but an initial microbiome is hard to change once established, and this is often cited as a reason for SynCom failure in more diverse habitats (Carlstrom et al. 2019, Shayanthan et al. 2022, Panchal et al. 2026). Given the presence of fungal contaminants, this could explain why isolate persistence was low and pioneers did not impact realized communities. The fungal contaminants could also have undermined the diversity treatments if the plants already had an established community that was difficult to colonize. In addition, low diversity taxa may have fulfilled the same functions as high diversity taxa even if they didn’t persist through the experiment. As priority and diversity treatments had little impact on plant performance and realized community composition, this supports that we should be focusing on creating microbial compositions that mimic natural but beneficial community structures.

SynComs affected plant height, root mass, and the root-to-shoot ratio independent of water availability, indicating that these fungal communities were not “rescuing” plants from stress. Moreover, the presence of individual persistent isolates, including two contaminants, reduced plant size regardless of drought treatment. *Stagonosporopsis, Didymella, Pyrenophora*, and contaminant taxa *Fusarium* and *Elsinoё,* are all known pathogens to a variety of crop plants. Our selection of taxa for SynComs was not based on their potential benefits per se, but these groups also include taxa that have been observed to act as commensals or beneficial endophytes (excluding *Elsinoё*) (Pham et al. 2025, Canare et al. 2026, Poveda et al. 2022, Liu et al. 2011, Zhang et al. 2026, Huang et al. 2021, Zhang et al. 2025, Wei et al. 2024, Manghwar et al. 2021, Hao et al. 2026). Therefore, the reduction in height and root mass and increase in leaf lesions in the presence of these taxa is possible, even though the taxa used to assemble synthetic communities were isolated from healthy adult wheat plants. These results are a good reminder that fungal behavior can be highly context dependent.

Microbial traits can sometimes be used to predict their effects on plant hosts (Giauque et al. 2018), which can be useful for generalizing across species. Here, however, the tested traits were not predictive of either realized communities or effects on host plant responses. It is possible that other traits might be more successful predictors. Additionally, culture-based traits may be poor predictors of behavior in the host. Improved predictions may require a more mechanistic approach, such as genomic or metabolic modeling focused on assembling SynComs with desired functional outcomes (Lin et al. 2026, Jourdain & Gu et al. 2025). For example, the combination of microbial genomics, microfluidic cultivation, and a machine learning framework was used to successfully create bacterial SynComs that degrade pollutants, including PFOS (perfluorooctane sulfonate), with high efficiency (De la Vega-Camarillo et al. 2026). Despite the narrow focus of SynCom development in waste or pollution cleanup, similar frameworks might be useful to develop plant-associated SynComs with certain outcomes, such as mitigating climate stressors, particularly if we can use their distributions as an initial screening tool. Looking to the future, the vast public repositories of microbiome data can be mined to more robustly select potential taxa for SynComs, and this approach could also support the development of more diverse cross-kingdom consortia for effective plant inoculations. In addition, we should leverage not only fungal data, while this simplifies experiments, but also include bacteria and yeasts as these microbes make up a large part of plant microbiomes and have important impacts on plant performance (Armanhi et al. 2021, Harman et al. 2021, Nimsi et al. 2024, Amprayn et al. 2012).

SynComs reduce microbial community complexity, allowing us to study the essential ecological processes that contribute to microbiome assembly and function (Shayanthan et al. 2022). However, as climate change progresses and plants encounter greater physiological stressors, ecological processes will change and therefore alter their relationship with microbes. Consideration of the establishment and persistence of these foliar communities will be important in improving current agricultural SynCom products’ performance (Compant et al. 2024). This study provides support for keystone/central taxa being essential players when developing persistent and beneficial SynComs. Therefore, when creating SynComs, especially with the goal of mitigating future stressors on plants, we should emulate field communities by studying microbial cooccurrence of keystones and their associated taxa.

## Acknowledgements

We are grateful to Haley Clayborne, Isabelle Webster, Angeleen Castillo, Breanna Lavalee, Xavious Allen, Rachel Hammer, Harry Ervin, Luciana Bachega, and Matthew Willman for assistance with lab work. The authors thank Christina Cowger, Marc Cubeta, Kevin Garcia, and Rytas Vilgalys for feedback on study design and research methodologies. Previous drafts of the manuscript were improved by comments from Rytas Vilgalys, Xavious Allen, and Elizabeth Paillan. This work was supported by funding from the Novo Nordisk Foundation (NNF19SA0059348) and by the Research Capacity Fund (HATCH; project award no. 7005451) from the U.S. Department of Agriculture’s National Institute of Food and Agriculture to CVH. In addition, BKA was supported by an NIH Tri Institutional Medical Mycology Training Fellowship (T32AI052080). The funders had no role in study design, data collection and interpretation, or the decision to submit the work for publication.

## Author Contributions

BKA and CVH designed the research. BKA established the experiment and collected the data. BKA analyzed the data with guidance from CVH. MD collected field samples, isolated fungal strains used in SynComs, prepared Illumina libraries, and ran the bioinformatic pipelines. BKA and CVH wrote the manuscript. BKA and CVH edited and revised the manuscript.

## Data Availability

Sequence data is available on NCBI (BioProject PRJNA1255089). Plant performance and fungal trait data are available on Github https://github.com/HawkesLab/Almeida-Hawkes_MATRIX and will be moved to Dryad before publication.

